# Reference-free Structural Variant Detection in Microbiomes via Long-read Coassembly Graphs

**DOI:** 10.1101/2024.01.25.577285

**Authors:** Kristen D. Curry, Feiqiao Brian Yu, Summer E. Vance, Santiago Segarra, Devaki Bhaya, Rayan Chikhi, Eduardo P.C. Rocha, Todd J. Treangen

## Abstract

Bacterial genome dynamics are vital for understanding the mechanisms underlying microbial adaptation, growth, and their broader impact on host phenotype. Structural variants (SVs), genomic alterations of 10 base pairs or more, play a pivotal role in driving evolutionary processes and maintaining genomic heterogeneity within bacterial populations. While SV detection in isolate genomes is relatively straightforward, metagenomes present broader challenges due to absence of clear reference genomes and presence of mixed strains. In response, our proposed method rhea, forgoes reference genomes and metagenome-assembled genomes (MAGs) by encompassing a single metagenome coassembly graph constructed from all samples in a series. The log fold change in graph coverage between subsequent samples is then calculated to call SVs that are thriving or declining throughout the series. We show rhea to outperform existing methods for SV and horizontal gene transfer (HGT) detection in two simulated mock metagenomes, which is particularly noticeable as the simulated reads diverge from reference genomes and an increase in strain diversity is incorporated. We additionally demonstrate use cases for rhea on series metagenomic data of environmental and fermented food microbiomes to detect specific sequence alterations between subsequent time and temperature samples, suggesting host advantage. Our innovative approach leverages raw read patterns rather than references or MAGs to include all sequencing reads in analysis, and thus provide versatility in studying SVs across diverse and poorly characterized microbial communities for more comprehensive insights into microbial genome dynamics.

## Introduction

Structural variants (SVs), loosely defined as genomic alterations that are 10 base pairs (bps) or longer (12), play an important role in driving both evolutionary adaptation and heterogeneity in bacterial genomes (31). Bacterial genome dynamics not only influence the ability for the bacteria to grow and adapt to changing environments (32), but can also impact the function of the microbial community as a whole and the phenotype of the host (11). In isolate genomics, the goal of SV detection is relatively straightforward: detect long genomic differences between a sequence and reference genome that can be classified as an insertion, deletion, inversion, duplication, translocation, or any combination of the prior (37). However, in metagenomics, when reference genomes may not be well-defined and a mixed population of similar strains may exist in the community, detection of SVs becomes more complex (37).

SV detection methods can be broadly categorized into three groups: mapping-driven, assembly-driven, and pattern-driven (Table 1) (37). In mapping-driven approaches, reads are directly aligned to an established reference genomes or pangenome of sequences, then unexpected mapping patterns identify SVs. In assembly-driven approaches, reads are first assembled into longer sequences (contigs), then aligned to another contig or reference to detect long scale differences. In pattern-driven approaches, SV patterns are pre-defined then search for in sequencing reads. Zeevi et al. developed a mapping-driven SV detection approach for metagenomic short reads to survey SVs associated with host disease risk factors in the human gut microbiome (42). The authors built a comprehensive database specifically for known microbes in the human gut microbiome and developed an “iterative coverage-based read assignment” (ICRA) algorithm to repeatedly adjust read assignments and establish alignments. Their SGV-Finder algorithm then scans the coverage of each reference genome for presence of regions with unexpectedly low (deletions) or high (duplications) coverage. While this method has been effective as a comprehensive search for SVs in the human gut microbiome correlating to expressed phenotypes (24), relying on a confident database of reference genomes is challenging for communities that have not been extensively characterized. This pipeline is additionally restricted to only deletions and duplications relative to reference genomes in the supplied database.

**Table 1.** Methods for SV detection in metagenomes, separated by types: mapping(M)-, assembly(A)-, and pattern(P)-driven. SV types abbreviations are as follows: Ins: insertion, Del: deletion, Dup: duplication, Inv: inversion, Trans: translocation, and CI: complex indel (defined here as an insertion and deletion at the same location). Input types are short reads (short), long reads (long), or metagenome-assembled genomes (MAG).

| Software | Type | Detected SV Types | Input |
| --- | --- | --- | --- |
| SVGFinder (42) | M | Ins, Dup | short |
| MetaSVs (23) | A | Ins, Del, Dup, Inv, Trans | MAG |
| MetaCHIP (35) | A | HGT Ins | MAG |
| PhaseFinder (16) | P | Inv | short |
| DIVE (1) | P | MGE Ins, MGE CI | short |
| Rhea | P | Ins, Del, Dup, CI | long |

To expand upon the types of SVs detected and leverage advantages of long read technologies, MetaSVs, an assembly-driven approach, was designed (23). In this pipeline, long and short reads combined help to confidently create and classify metagenome-assembled genomes (MAGs). Each MAG is then evaluated independently through whole-genome alignment to a reference MAG or genome with the SV detection tool MUM & Co (29). Chen et al. utilized MetaSVs to expand upon characterized SVs in the human gut (notably insertions and inversions) and demonstrates the value in incorporating long reads for SV detection (9). However, this assembly-driven method is still highly dependent on a reference database, as it is the taxonomic reference-driven classifications that determine which MAGs get compared to which references. Additionally, unique MAGs are often not created for subtle SV differences (18), especially in microbial communities where similar strains are present (14).

MetaCHIP is another MAG-based approach for the slightly different goal of detecting recent horizontal gene transfer (HGT) events within a metagenome (36). In an HGT event, genetic material is exchanged between organisms (28), resulting in an insertion SV for the recipient microbe. MetaCHIP effectively evaluates each MAG in the community for a gene sequence that has more BLASTN (2) hits to genes in a different MAG than its own. This algorithm, however, can only detect insertion genes that are highly similar to another MAG, which resulted in simulation results declining at 25% mutation rate between donor and recipient.

To entirely avoid reference genomes and MAG creation, two pattern-driven methods have been developed. PhaseFinder (16) was created for detection of inversions in bacterial genomes from genomic or metagenomic data, by detecting regions flanked by inverted repeats where sequencing reads support both orientations. DIVE (1) was developed in 2023 to identify sequences surrounding genetic diversification such as transposable elements, within MGE variability hotspots, or CRISPR repeats, by detecting constant k-mers with diverse flanking sequences to define MGE bounding sequences and transposon arms. While both these methods show how patterns in raw read can be used to eliminate reference genomes and MAGs, they are limited to only these specific patterns.

Rhea takes a different approach to detect SV patterns within a microbial community. It constructs a coassembly graph from all metagenomes in a series that are expected to have similar communities (i.e. longitudinal time series or cross-sectional studies where a significant portion of the strains are shared across samples). Regions of the graph indicative of SVs are then highlighted, as previously explored for characterization of genome variants (27; 13). The log fold change in graph coverage between consecutive steps in the series is then used to reduce false SV calls made from assembly error, account for shifting levels of microbe relative abundance, and ultimately permit SV detection in understudied and complex microbial environments. Recent work utilizes coassembly graphs for metagenomes to decompose strain diversity into haplotypes (30), but to the best of our knowledge, this is the first time coassembly graph patterns have been used for automated detection of SVs in a metagenome series.

## Methods

### Rhea method

Rhea takes as input a series of long-read metagenomic sequences, expected to be taken from the same source at different time points or some other step-wise metadata separation. A single metagenome assembly graph is constructed by combining all provided samples, then each sample is separately aligned back to the graph. Change in graph coverage between subsequent samples and the graph structure are used to call SVs (Figure 1).

**Fig. 1.**
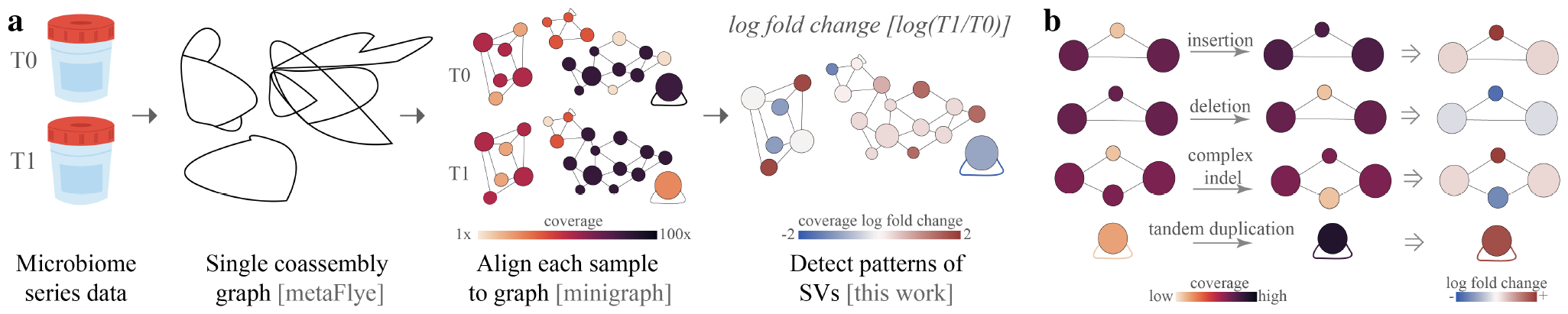
(a) To utilize rhea, first, microbiome series data must be collected and long whole genome sequencing reads generated. Then, within rhea, a coassembly graph of all reads in the series is created with metaFlye. Reads from each sample are then separately aligned to the coassembly graph with minigraph. Rhea evaluates log fold change in coverage between series steps for SV-specific patterns in the assembly graph to detect structural variants between steps. (b) Assembly graph patterns detected in rhea, which indicate potential insertions, deletions, complex indels, and tandem duplicates. Insertions and deletions are detected by observing a triangle where one node has a significantly higher (insertion) or lower (deletion) log fold change. Complex indels are noted by a square with one or two outliers; in the case of two outliers, the two outliers must be of opposing sides of the median and not have an edge between them. Tandem duplicates are detected by a log fold change of a self-loop edge coverage greater than 1.

### SV definitions

Four types of SVs are detected in rhea: insertions, deletions, tandem duplications (37), and complex indels (41; 33). An insertion here is a sequence that has been integrated in increasing abundance between subsequent steps in the series. A deletion is the opposite, a subsequence that is declining. A tandem duplication is a gene sequence that has been repeated, directly one after another, in increasing presence. A complex indel as a sequence that has drastically changed between subsequent steps, showing the signature of a deletion and insertion at the same location. In this pipeline, SV detection equates to an increase in abundance of the SV, rather than simply a novel appearance, and therefore suggests a provided advantage for the host microbe or the community.

### Graph construction and coverage calculations

A single coassembly graph for the series with *N* samples is constructed by combining all reads from all samples into one metaFlye run (19), with `--keep-haplotypes` parameter set to true to maintain strain variations. After the graph is constructed, each sample is separately aligned back to the graph with minigraph (22). An undirected graph is then built mimicking the structure of the metaFlye assembly graph where a single node is drawn for each complementary pair, as seen in the assembly graph visualization software Bandage “single” option (38). This graph is defined as *G* = (*V, E*) with a set of *k* nodes *V* = {*v*_1_, *v*_2_.., *v*_*k*_} and a set of edges *E*. Each edge (*e*_*i,j*_) is then given a weight equal to the number of edges that appear between nodes *i* and *j* in the metaFlye assembly graph, given there exist at least one edge between *i* and *j* in the assembly graph. Each edge (*e*_*i,j*_) thus denotes the existence of overlap reads that expand directly from *v*_*i*_ to *v*_*j*_ (or from *v*_*j*_ to *v*_*i*_) without gaps, in either direction (forward or reverse) for the sequences in *i* and *j*. Minigraph alignments are then used to calculate node and edge coverage for each step in the series. Node coverage is calculated as the average coverage per base pair within the node, calculated by summing the coverage for each base pair divided by the total number of base pairs in the node. To account for error, all nodes with coverage less than 1, are set to a coverage of 1. Node coverage is then normalized for the entire series, by first calculating the median total base pairs *m* across samples in the series, then establishing a multiplier for each sample *n* = 0..*N* as *bp*_*n*_*/m*, where *bp*_*n*_ is the number of base pairs in sample *n*. This multiplier for each step is applied to all node coverage for each *n* = 0..*N*. Edge coverage for each edge *e*_*i,j*_ at each step *n* in the series is counted as the number of occurrence a read path covers directly from *i* to *j* or *j* to *i* in the read-graph alignment for step *n*. Each node in our undirected assembly graph then holds a vector of log fold change in coverage between subsequent steps in the series, calculated for each node *i* as 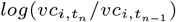, where *vc*_*i,t*_ is the coverage of node *i* at step *n* in the series for all steps *n* = 1…*N*. A log fold change vector is also assigned to each edge (*i, j*), defined as 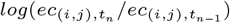, where *ec*_*i,t*_ is the coverage of edge *e*_*i,j*_ at step *n* in the series for all steps *n* = 1..*N*. The log fold change vectors are then used in the next step to detect SVs and account for assembly error and changes in genome relative abundance between subsequent samples.

### Detected SV graph patterns

Rhea utilizes the graph structure, edge weights, and the log fold change coverage vectors to call SVs between each pair of consecutive samples in the series. For insertions and deletions, each triangle is searched for the pattern of two similar log fold change values and one that is significantly different for each step. This is completed by: calculating the median and standard deviation between the three log fold change, then labeling any node with a value that is more than one standard deviation away from the median as an outlier. If the triangle contains exactly one outlier, then an insertion or deletion is called, depending on if the outlier value is lower (deletion) or higher (insertion) than the median. Median is used here rather than mean to provide robustness against extreme outliers. For example, in the case of an extreme outlier due to a deletion from a thriving member in the community, the mean would be skewed and thus could call all three nodes an outlier; whereas the median would take the value of one of the non-deletion nodes and thus, given the two non-deleted nodes carry a similar value, only the deletion would be an outlier. A similar process is conducted to search for complex indels. Here, each square (cycle of length 4) in the graph is searched for outliers. If the square either has a single outlier or two outliers that do not have an edge between them (opposites in the square) and one is greater than the median while the other is smaller, a complex indel is called. A tandem duplicate can be called under two different scenarios. The first, a self-duplicate, shown by an edge log fold change of any self-loop edge greater than 1 for any subsequent steps in the series. The second is the situation where the duplicate produces a second node containing a nearly duplicate sequence and loops between two nodes. This is detected by searching all edges with weight *w ≥* 2 for a log fold change edge weight greater than 1. If these criteria are met, the node with the greater log fold change coverage between the two is then called a tandem duplication if it has not been called for another SV at the specified step.

## Experiments

### Simulated HGT events

Rhea was compared to the metagenome HGT detection tool MetaCHIP by simulating long reads from the simulated HGT events completed in the HgtSIM manuscript (35). For this community, 10 strains within class *Alphaproteobacteria* and 10 strains within class *Betaproteobacteria* were selected. 1 gene was selected from each *Alphaproteobacteria*, mutated with rate *m*, and inserted randomly into each *Betaproteobacteria*. This resulted in a total of 100 HGT events for the community (Fig 2a). Three long read metagenomic datasets of 500,000 reads were simulated from these reference genomes with NanoSim (40) v3.1.0 with default parameters: a pre-transfer community (*T0*) of the 20 reference genomes in equal abundance, and two separate post-transfer communities with mutation rate *m* = 0 and *m* = 30 (*T* 1_*m*0_, *T* 1_*m*30_), which include the 10 original *Alphaproteobacteria* and the 10 HGT-inserted *Betaproteobacteria* references in varying abundances (Fig 2b). These varying abundances were established by randomly selecting relative quantity between 1 and 5 for each of the species as input into the NanoSim abundance text file. MetaCHIP v.1.10.12 was run with GTDB-Tk (8) v2.2.6 with taxonomy release 207 and `-r` set to class (c). Rhea v1.0 was run with default parameters, metaFlye v2.9.3, and minigraph v0.20. Simulated HGT insertions were mapped against reported HGT sequences for both methods using minimap2 (21) v2.24 with default parameters; each HGT insertion sequence was marked as detected if the sequence had a hit to a reported HGT insertion.

**Fig. 2.**
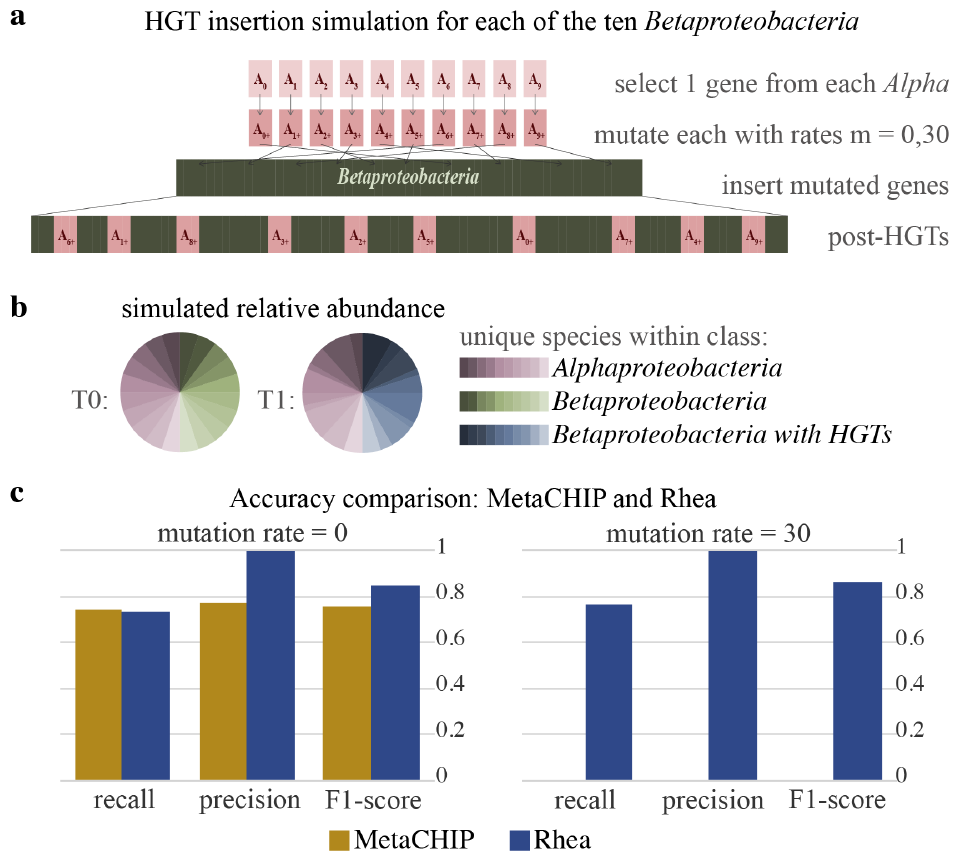
(a) HGT simulation process completed in the HgtSIM publication (35). One gene is randomly selected from each of the 10 *Alphaproteobacteria* species, mutated with rate *m*, then inserted into each *Betaproteobacteria*. Mutations rates *m* = 0 and *m* = 30 are included in this study. (b) Simulated relative abundances for time points *T0* and *T1. T0* is a simulation of the 20 reference genomes in equal abundance; *T1* is simulated from the 10 original *Alphaproteobacteria* species and the 10 mutated *Betaproteobacteria* species in varying abundances (c) Precision, recall, and F1-score for MetaCHIP (36) and rhea detected insertions for the mock community with mutation rates 0 and 30. Time point *T1* is used for MetaCHIP results; change from *T0* to *T1* is used for rhea.

### Simulated SVs

To evaluate the accuracy of rhea for detection of SV types insertion, deletion, complex indel, and tandem duplication in comparison with a MAG-based workflow, variants of each of the 10 microbes in the ZymoBIOMICS Microbial Community Standard were generated. SURVIVOR (15) v1.0.7 was used to randomly create 20 indels (insertions or deletions) and 10 tandem duplicates of length 500-2000 base pairs, with homozygous ratio=0.5 and Number haploid=1 in the parameters file, for each of the 10 reference genomes independently. Then a custom script introduced 10 random complex indels of the same length range into each of the variant strains. The custom script randomly selected a location along the genome, then performed a deletion and a random insertion, each within the prescribed length range. Two long read metagenomic datasets of roughly 500,000 reads were simulated from these reference genomes with NanoSim: a pre-transfer community (*T0*) of the original references in their provided relative abundances and a post-transfer community (*T1*), which includes only the variant strain for half of the species and equal abundance of variant and original strains for the other half (Fig 3a). For our MAG workflow, reads were assembled with metaFlye (19) with --keep-haplotypes set to true, contigs were binned with MetaBat (17) v2.15 with default parameters, and bins were classified with GTDB-Tk. Bins with the same classification in both simulated samples were analyzed for SVs with MUM & Co (29) v3.8 with the known reference genome length for parameter `-g`. Simulated SV sequences were mapped against reported SV sequences for both methods using minimap2. Each simulated SV was marked as detected if the sequence had a hit to a reported SV sequence with the correct SV type. Since MUM & Co does not call complex indels, we considered these correct if both the deletion sequence and the insertion sequence were returned.

**Fig. 3.**
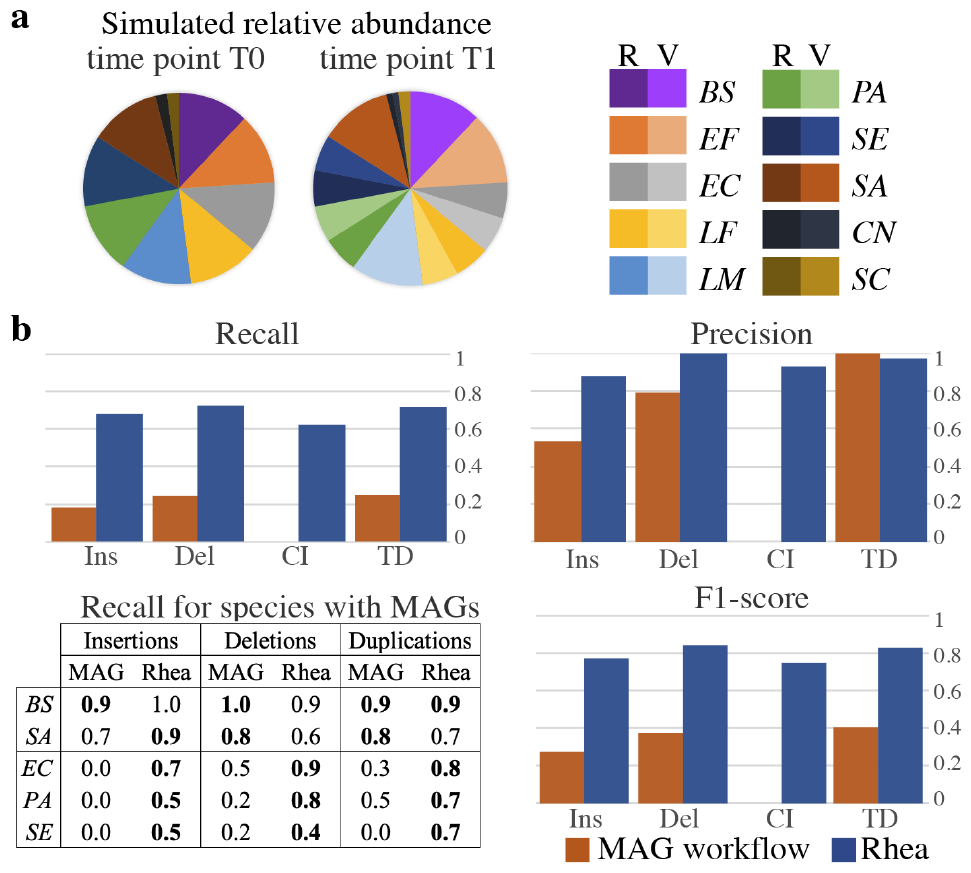
(a) Relative abundance of long reads for two simulated time points (*T0, T1*) for our ZymoBIOMICS community. Each of the 10 microbes were randomly given 20 indels, 10 tandem duplications, and 10 long complex indels to create a variant strain (15). *T0* contains only the original references (R); *T1* introduces the variants (V), where half the species have variants in equal abundance to their original reference *[Escherichia coli (EC), Lactobacillus fermentum (LF), Pseudomonas aeruginosa (PA), Salmonella enterica (SE), Cryptococcus neoformans (CN)]*, and half the species are dominated by their variants *[Bacillus subtilis (BS), Enterococcus faecalis (EF), Listeria monocytogenes (LM), Staphylococcus aureus (SA), Saccharomyces cerevisiae (SC)]*. (b) Complete recall, precision, and F1-score for each of the SV types (Ins: insertion, Del: deletion, CI: complex indel, TD: tandem duplication) for both workflows (bar plots) and recall on a subset of 5 species (table). For the MAG workflow, MAGs were curated for *T0* and *T1* separately. Then, Mum & Co called SVs between *T0* and *T1* MAGs of matching taxonomic classification. The 5 species selected for the table are the 5 species with a classified MAG at both time points. The top portion (*BS, SA*) show the species where the variant dominates in *T1* ; whereas both the variant and the original reference are present in *T1* for the bottom portion (*EC, PA, SE*). The better recall is in **bold** for each comparison.

### Cheese rind ripening

To evaluate rhea on a real microbiome, PacBio HiFi metagenomic reads from cheese rinds throughout ripening were taken from a previous study (34). One rhea run for “Cheese C” was completed with the 5 corresponding samples in temporal order and parameter --type set to pacbio-hifi. The assembly graph connected component that showed interesting evolutionary patterns was classified with GTDB-Tk (8) “classify-wf” with default parameters, and is referred to rhea 5 as the *Halomonas* subgraph per this taxonomic classification. Mobile genetic element (MGE) contigs and putative hosts were established in the original publication utilizing Hi-C sequencing technology, overlap read coverage, and the viralAssociatePipeline (6). To determine which of these contigs showed signatures in our *Halomonas* subgraph, BLAST (2) was run for all MGE contigs with a putative host, against the extracted *Halomonas* subgraph sequences as reference with default parameters. MGE contigs were considered to have their signatures present in the graph if a hit with query coverage *>* 5% was reported. One subsection of the *Halomonas* subgraph was selected for further investigate as it showed a change in dominating graph path over time. Nodes within this path were characterized with SeqScreen-Nano (3) v4.1 with default parameters and provided SeqScreen databases v21.4.

### Hot spring microbial mat sequencing

Microbial mat plugs were extracted from Mushroom Spring, Yellowstone National Park, USA on July 30, 2009 across a series of temperatures: 50^*°*^C, 55^*°*^C, 60^*°*^C, 65^*°*^C. DNA was quantified using the Qubit 3.0 Fluorometric Quantitation dsDNA High Sensitivity kit (ThermoFisher Scientific, Waltham, MA, USA) and stored for future use at -80^*°*^C. DNA extractions were analyzed using the Genomic DNA ScreenTape Analysis kit on the 4150 TapeStation System (both from Agilent, Santa Clara, CA, USA). Size selection using AMPure XP beads (Beckman Coulter, San Jose, CA, USA) increased DNA fragment length from a mean of 2kb up to 6kb with high recovery of DNA. Size selected DNA was prepped for sequencing using the Oxford Nanopore Technologies (ONT) 1D Genomic DNA by Ligation library preparation kit (SQK-LSK109, Oxford Nanopore Technologies, Oxford, UK). Libraries were then sequenced using the ONT MinION sequencer using one FLO-MIN106D R9 Version Rev D flow cell per temperature sample. Sequencing was run on a MacBook Pro (model A1502, Apple) using ONT’s MinKNOW software. Automatic basecalling through this software was turned off. Sequencing runs lasted between 24-44 hours. Basecalling was completed using the ONT software Guppy (https://github.com/nanoporetech/pyguppyclient.git) with default parameters.

### Hot spring microbial mat analysis

Rhea was run on Oxford Nanopore Technologies (ONT) reads from a hot spring microbial mat for 4 unique temperatures (see above) to asses an environmental microbiome with a high-level of complex microbial interactions (5; 26). Basecalled sequences were listed in order of increasing temperature with the `--collapse` parameter set to true. MAGs were also curated for reads from the 60^*°*^*C* sample by metaFlye assembly with `--keep-haplotypes` set to true and contigs binned with MetaBat 2 (17). Each read was then aligned back to the set of MAGs with minimap2 with default parameters. Reads with an alignment to a MAG contig of *>* 80% of length were considered to be included in MAGs, mimicking the pipeline of a previous manuscript (4). Kraken 2 (39) v2.1.1 was additionally run with the Kraken 2 default parameters and RefSeq indexes released on May 17, 2021 for all raw reads in this sample.

## Results

### Simulated HGT insertions

Two simulation experiments were conducted with a community of strains within *Alphaproteobacteria* and *Betaproteobacteria* classes to evaluate HGT detection accuracy: one with mutation rates *m* = 0 and the other with *m* = 30. For the HGT insertions with *m* = 0, rhea delivered comparable recall to MetaCHIP (0.73 to 0.74) and improved precision (1.0 to 0.77) (Fig 2c). The only non-insertion SV that rhea called was a single complex indel, which was due to two insertions sequences in close genomic proximity. Given the two inserted sequences were still detected as sequences of increasing abundance, this was still considered this an accurate call. Although results for MetaCHIP and rhea for *m* = 0 were relatively similar, a large discrepancy was observed for mutation rate *m* = 30. Here, the accuracy for rhea stays consistent to that of no mutations (0.76 recall and 1.0 precision), yet MetaCHIP is not able to detect any of the HGT insertions. This caveat is also highlighted in the MetaCHIP manuscript; the inserted sequence is required to be present in another MAG (putative donor) in the community for MetaCHIP to be able to detect the HGT insertion. Additionally, MetaCHIP returned a total of 13 false positive insertions, while rhea did not report any false positives.

### Simulated structural variants

A single simulated experiment was conducted to evaluate rhea in comparison to a MAG-based workflow for a variety of SVs. This experiment contained two mock time points (*T0* and *T1*), where *T0* contains only the references in the ZymoBIOMICS Microbial Community Standard and *T1* contains a mix of original references and simulated variants. For the 400 simulated SVs, rhea greatly outperformed the MAG workflow in terms of recall (Fig 3). While rhea detected 71, 68, 63, and 72 of the simulated insertions, deletions, complex indels, and tandem duplications respectively, the MAG workflow only identified 19, 23, 0, and 25, respectively. This discrepancy was largely due to the inability to curate independent MAGs for low abundant species and SV distinctions.

MAGs were classified for 5 of the 10 species at both *T0* and *T1*, limiting the MAG-based workflow to only attempt to call SVs for these species. Of the 5 species, 2 (*B. subtilis, S. aureus*) were from species where the SV-containing strain dominated in sample *T1*, while 3 (*E. coli, P. aeruginosa, S. enterica*) contained both the original and the SV-containing strains in *T1*. Accuracy results between rhea and MAG pipelines proved comparable for insertions, deletions, and tandem duplicates when only the SV-strain was present in post-transfer sample *T1*. However, when both the original and SV-strains were present, only one MAG was curated for the species, leaving many of the SV graph nodes unbinned and thus impossible to detect. Since the SV caller used in the MAG workflow does not call complex indels, we considered a complex indel to be detected if both the insertion and deletion for the complex indel was reported; however, this was not the case for any of the 50. For the two low abundant fungi present in only 2% relative abundance, MAGs were not created at either time point, while rhea was able to detect SVs for these species with similar recall to the more abundant bacteria. Even with the reduced potential to call SVs with a MAG-based workflow, this process resulted in 21 false positive SVs call while rhea only elicit 17.

Of the 125 SVs that were not detected by rhea, roughly 50% were not detected in the assembly graph, roughly 40% were in the graph but resolved into longer nodes rather than partaking in SV graph patterns, and the remaining 10% were called as the wrong SV type.

### Cheese ripening temporal series

To demonstrate rhea’s ability to extract interesting microbial evolutionary patterns within a microbiome over time, PacBio HiFi metagenomic sequences taken from a cheese rind over the course of ripening were used as input (34). A total of 5 samples were included from sampling weeks 2, 3, 4, 9, and 13, creating 4 pairs of change (*C1-4*). Evaluating the assembly graph coverage visuals produced by rhea and Bandage (38), one connected component stood out for displaying significant graph complexity and diversity in coverage, implying a disproportionately large number of SVs. Rhea SV results indicated roughly 20% of SVs in the community to be contained in this subgraph (Fig 4a). This connected component was then classified by GTDB-Tk under genus *Halomonas* and further exploration was pursued.

**Fig. 4.**
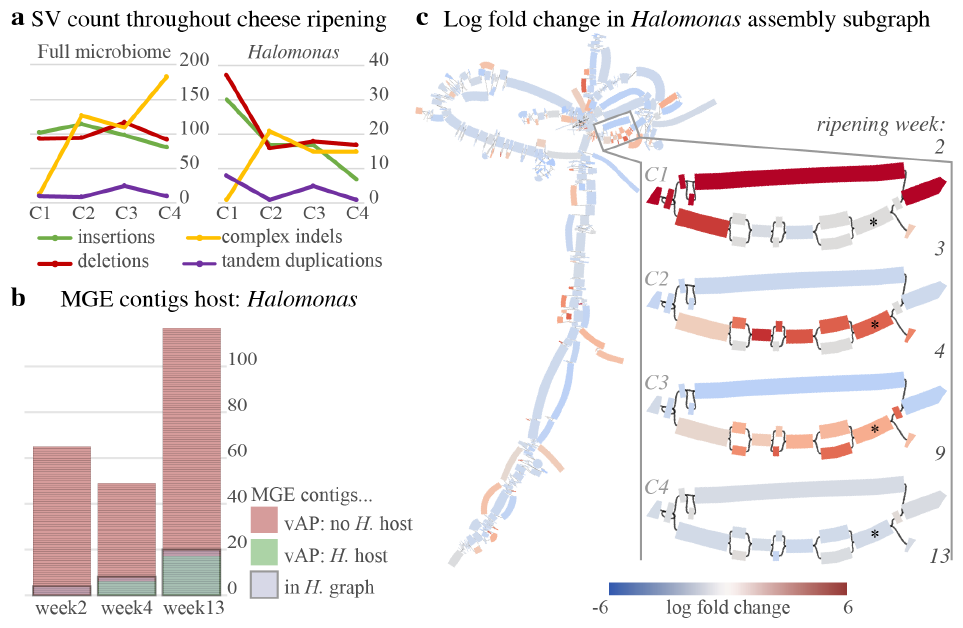
(a) SV counts detected by rhea for pairs of subsequent samples throughout cheese ripening (*C1-4*) for the entire community and exclusively the extracted *Halomonas* subgraph. (b) Previously established MGE contigs for 3 selected time points, described as either with (green) or without (red) *Halomonas* host by viralAssociationPipeline (vAP) per original publication’s findings. Grey boxes signify the MGE contigs that had a BLAST hit of *>* 5% query coverage to our *Halomonas* subgraph. (c) Rhea and Bandage generated visual for the log fold change in coverage for the *Halomonas* subgraph. Left shows the complete *Halomonas* subgraph between weeks 4 and 9 (*C3*), selected for showing a general decrease in abundance yet an increase in abundance for several subsequences. Right zooms in on a small portion of the subgraph containing an interesting evolutionary pattern, where the log fold change in coverage graph is shown for each pair of subsequent time points (*C1-4*). The graph node marked with a *∗* indicates the node containing the predicted type I restriction-modification system.

First, the ability for viral and plasmid mobile genetic elements (MGEs) to show signatures in the *Halomonas* subgraph was evaluated. In the original publication for the cheese samples, MGE contigs and putative hosts were established via Hi-C sequencing technology and overlap read coverage with the viralAssociatePipeline (6) for sampling weeks 2, 4, and 13. Their results showed *Halomonas* to be host for 0, 6, and 17 MGE contigs, respectively. A BLAST (2) comparison of all MGE contigs against the *Halomonas* subgraph, showed all putative *Halomonas* MGE contigs to display signatures in our *Halomonas* subgraph (hit with more than 5% query coverage), despite previous host connections being defined via Hi-C sequencing and our graph being constructed solely on long-read sequences. An additional 4, 2, and 3 MGE contigs showed signature in the *Halomonas* subgraph without having a previous description of a *Halomonas* host for the time point for each of the 3 included sampling weeks respectively (Fig 4b), which may be false positives or novel host discovery. Finally, one striking section of the *Halomonas* subgraph was selected for gene function analysis (Fig 4c). Here, a newly emerged path (displayed lower option) shows an increase in coverage over time up until stabilizing by week 9, suggesting an evolutionary advantage over the alternative path (top option). Gene function predictions returned by SeqScreen (3) showed the newly dominating path to contain a type I restriction-modification system that was not expressed in the alternative sequence. This suggests an evolutionary advantage due to phage protection in the *Halomonas* strains, which is unsurprising given the increasing number of phage interactions detected throughout ripening for *Halomonas*. Exploratory analysis here demonstrates a novel approach produced by rhea to extract genomic subsequences that suggest an evolutionary advantage, gain insight into MGE hosts, and infer microbial interactions.

### Hot spring microbial mat temperature series

Lastly, to assess an environmental sample with complex interactions, rhea was run on a temperature series of samples taken from the Mushroom Spring microbial mat in Yellowstone National Park, USA. Samples were collected from 4 different portions of the mat with temperatures 50^*°*^C, 55^*°*^C, 60^*°*^C, and 65^*°*^C. Rhea detected SVs between subsequent temperature increases (Table 2). An extraordinarily large number of SVs were detected in the hot spring microbial mat, averaging 8.9 million per consecutive pair, as opposed to an average of 317 per pair in the cheese microbiome. The vast quantity of SVs is particularly noticeable for complex indels, as counts for this type was observed to be over an order of magnitude greater than the other SV types observed. The number of detected complex indels increased with the first two temperature increases (over 8 million and 22 million, respectively), but then fewer are detected with the last temperature increase (over 3 million). While this decrease implies more stability at these higher temperatures, a closer look at the coassembly graph and alignments could confirm this pattern is true signal rather than a result of decreased average read length in the 65^*°*^C sample. Previous research closely analysed two *Synechococcus* isolates from these mats and showed a large number of diverse insertion sequence (IS) activity occurring within the two strains (26). Our findings suggest there is far more transposon and gene exchange occurring in microbial mats that has yet to discovered, and likely many uncharacterized novel bacterial strains. Further research is needed to confirm these suspicions and additionally detect the gene functions for the thriving SVs to give insight into evolutionary drivers for these extremophiles.

**Table 2.** Sample and SV statistics for hot spring microbial mat temperature series. SV counts shown represent the number of SV detected between the sample listed in the row and the previous row. SV types abbreviations are as follows: Ins: insertion, Del: deletion, TD: tandem duplication, and CI: complex indel.

| Sample | Reads<br>(million) | Bps<br>(billion) | Ins<br>(10<sup>3</sup>) | Del<br>(10<sup>3</sup>) | TD<br>(10<sup>3</sup>) | CI<br>(10<sup>3</sup>) |
| --- | --- | --- | --- | --- | --- | --- |
| 50°C | 3.6 | 7.5 |  |  |  |  |
| 55°C | 2.4 | 6.2 | 224 | 232 | 0.21 | 8616 |
| 60°C | 3.4 | 7.7 | 220 | 239 | 0.38 | 22217 |
| 65°C | 2.9 | 3.7 | 212 | 242 | 0.19 | 3611 |

**Table 3.** Computational usages for rhea experiments.

| study | reads<br>(million) | base pairs<br>(billion) | User+sys<br>time (h) | RAM<br>(GB) |
| --- | --- | --- | --- | --- |
| HgtSIM (m0) | 1.0 | 4.0 | 13 | 26 |
| HgtSIM (m30) | 1.0 | 4.0 | 13 | 26 |
| ZymoBIOMICS | 1.0 | 4.0 | 13 | 26 |
| Cheese | 1.8 | 23.1 | 154 | 47 |

One sample (60^*°*^C) was selected to assess read inclusion rate of alternative workflows for this community rife in unknown microbes. To evaluate a reference-based taxonomic classification method, reads were classified by Kraken2 with default database, where 42% of the reads were left unclassified. To evaluate a MAG creation workflow, MAGs were created with MetaFlye contigs and MetaBat2 binning, where roughly 30% of raw reads did not map to a binned contig. Use of rhea allowed for the inclusion of all sequenced reads to distinguish subsequences and genomic context specific to high temperature environments and give insight into the evolutionary history of these active and uncharacterized microbes within hot spring microbal mats.

### Computational usage

All software analysis was completed on a Ubuntu 22.04 LTS system with 15 threads. The /usr/bin/time command was used to gather time and memory statistics. Reported CPU (central processing unit) time was calculated by summing the user and the system time; RAM (random access memory) requirements were determined using the maximum resident set size.

## Discussion

Here we present rhea, a novel method for detecting structural variants (SVs) between consecutive samples in long-read metagenome series data. Rhea leverages sequence information from the entire metagenomic community and avoids need for a reference database or MAG creation by analyzing structural motifs and change in alignment coverage on a combined coassembly graph. This permits SV detection for intra-species variations, low abundance genomes, and novel organisms. Our simulated results of recent HGT events and SVs in two mock communities show rhea to outperform existing methods. Use of rhea on a cheese rind microbiome with samples taken throughout ripening allowed us to infer MGE hosts that align with Hi-C sequencing and additionally suggest recently transferred genes with a suspected evolutionary advantage for the host. Use of rhea for a varying temperature series of samples from a hot spring microbial mat allowed us to include reads that would likely have been removed in alternative workflows, as strain-level diversity prevents sequences from being incorporated in MAGs and lack of isolate reference genomes prevent use of reference-based approaches. While extracting evolutionary insights from this complex community still provides a significant challenge, rhea introduces a first step in logically parsing these metagenomic sequences.

Methods for identifying significant changes throughout a metagenome series is an active area of research (43). Currently, a common approach is to first simplify each metagenome into a profile that can be logically aligned and compared, such as taxonomic classification relative abundance, gene function presence, and counts of short sub-sequences (k-mers) (10). Yet, each of these strategies either oversimplifies potentially important sequences of microbial communities or is biased by a reference database (20; 25). Rhea results contain input data for the interactive visual software package Bandage (38), for exploration of changes in graph coverage throughout a metagenome series. This tool provides researchers with an efficient method to investigate sequence-level fluctuations while maintaining genome context, to ultimately extract sequences of interest (Fig 4c). It is important to note that metagenomic sequences simply provide a snapshot of the microbial community at the time of sampling, and thus oscillating fluctuations that take place between samples may not be detected.

Currently, rhea is only able to detect insertion, deletion, tandem duplication, and complex indel SV types between two metagenomes of similar microbes. The method could theoretically be expanded to inversions and translocations, however, we anticipate the need to maintain node directionality (whether the sequence is read forward or reverse) in the evaluated coassembly graph. Rhea could also be expanded to detect more complex patterns of multiple overlapping SVs or short read sequences, but further experimentation is required. Rhea has so far only been evaluated for SV detection over the course of microbiome series data. The idea of constructing a coassembly graph and comparing the coverage between samples could be expanded beyond series data and used for different types of studies, such as cohort comparison analyses and MGE host detection. As the number of reads included in the study increases, methods of downsampling sequences to generate the graph or an alternate graph construction methods could be considered. Alternative graphs, could also be explored in attempt to improve sensitivity for SV detection, given that results in our mock ZymoBIOMICS community still collapsed nearly a quarter of simulated SVs. However, alternative graph structures could also create two unique connected components for microbes that have undergone significant structural variations, which would prevent the current detection algorithm within rhea to call such SVs. Further analysis could help determine at which diversity levels SVs are collapsed in a single node or separated into unique connected components, to provide genome similarity requirement guidelines for SV detection capability within rhea.

In lieu of metagenome-specific methods, metagenomes are often construed to fit methods and models developed for genome analyses. Yet this simplification overlooks inherent complexities of dynamic and interdependent microbial ecosystems (7). By viewing these communities holistically and acknowledging their intricate interplay and co-evolution, we can discover nuanced patterns, novel relationships, and a deeper understanding of the collective behaviors throughout the community. Developed to embody this ideology, rhea is a novel technique to pinpoint microbial heterogeneity and evolution by capturing the full essence of these diverse and interconnected ecosystems.

## Data availability

Rhea and all associate code are available on GitHub (https://github.com/treangenlab/rhea). Scripts, simulations, complete results, and hot spring long reads are available on OSF under project FVHW8.

## Author contributions statement

K.D.C. derived the concept, developed the software, completed experiments and wrote the manuscript. S.E.V. conducted sequencing. B.Y., D.B., S.S., R.C., E.P.C.C. and T.J.T. contributed to method development. All authors read, revised and approved the manuscript.

## Acknowledgments

K.D.C. was supported in part by Ken Kennedy Institute Computational Science and Engineering Graduate Recruiting Fellowship, Rice University Wagoner Foreign Study Scholarship, and the Chateaubriand Fellowship. T.J.T. was supported in part by NIH grant P01-AI152999 from the National Institute of Allergy and Infectious Diseases (NIAID) and NSF CAREER award IIS-2239114 (PI Treangen). K.D.C., S.S., and T.J.T. were supported by the NSF MIM Universal Rules of Live (URoL) grant (EF-2126387, PI Treangen). This project has received funding from the European Union’s Horizon 2020 research and innovation programme under Marie Sk-lodowska-Curie grant agreement Nos. 872539 and 956229 (R.C.). R.C. was supported by ANR Transipedia, SeqDigger, Inception and PRAIRIE grants (ANR-18-CE45-0020, ANR-19-CE45-0008,PIA/ANR16-CONV-0005, ANR-19-P3IA-0001). D.B. acknowledges support from a BBSRC-NSF/BIO collaborative research grant (award number 1921429), NSF Emerging Frontiers grant (award number 2125965); Joint Genome Institute Community Sequencing Project # 503441 and # 509352, and the Carnegie Institution for Science. The work conducted by the U.S. Department of Energy Joint Genome Institute, a DOE Office of Science User Facility, is supported by the Office of Science of the U.S. Department of Energy. S.E.V. is supported by NSF GRFP (fellow ID 2023333162).

